# *Schistocephalus* parasite infection alters sticklebacks’ movement ability and thereby shapes social interactions

**DOI:** 10.1101/849737

**Authors:** Jolle W. Jolles, Geoffrey P. F. Mazué, Jacob Davidson, Jasminca Behrmann-Godel, Iain D. Couzin

**Author notes:** Correspondence should be addressed to: Jolle W. Jolles.

## Abstract

Parasitism is ubiquitous in the animal kingdom. Although many fundamental aspects of host-parasite relationships have been unravelled, few studies have systematically investigated how parasites affect organismal movement. Here we combine behavioural experiments of *Schistocephalus solidus* infected sticklebacks with individual-based simulations to understand how parasitism affects individual movement ability and how this in turn influences social interaction patterns. Detailed movement tracking revealed that infected fish swam slower, accelerated slower, turned more slowly, and tended to be more predictable in their movements than did non-infected fish. Importantly, the strength of these effects increased with increasing parasite load (% of body weight), with the behaviour of more heavily infected fish being more impaired. When grouped, pairs of infected fish moved more slowly, were less cohesive, less aligned, and less coordinated than healthy pairs. Mixed pairs exhibited intermediate behaviours and were primarily led by the non-infected fish. These social patterns emerged naturally in model simulations of self-organised groups composed of individuals with different speeds and turning tendency, consistent with changes in mobility and manoeuvrability due to infection. Together, our results demonstrate how infection with a complex life cycle parasite affects the movement ability of individuals and how this in turn shapes social interactions, providing important mechanistic insights into the effects of parasites on host movement dynamics.

## Introduction

Parasitism is ubiquitous across the animal kingdom, with parasites often exerting considerable influence on their hosts, consuming energy, and inducing morphological, physiological, and behavioural changes [1–3]. Parasites have been best studied in terms of their ecological and evolutionary effects on hosts [1,4], but may also have large effects on host behaviour [2]. In particular, by changing their morphology and physiology, parasites may strongly alter the movement dynamics of their host [2,5], such as by reducing the energy available to allocate to movement or by compromising their movement capacity or mobility. So far, few studies have systematically investigated the mechanistic basis of such potential behavioural modifications [2,5] or considered the repercussions they may have for social interactions and group-level patterns via self-organising effects [6,7].

A model system for studying host-parasite relationships with well-documented parasite effects on host behaviour is that of three-spined sticklebacks *(Gasterosteus aculeatus)* infected with the plerocercoid larvae of the cestode *Schistocephalus solidus* [8,9], a complex life cycle parasite (i.e. not directly transmittable). The plerocercoids live in the body cavity where they can grow very rapidly, sometimes weighing as much as their host [10], and grossly distending their hosts’ body [11–13] (see Figure 1). *S. solidus* makes significant demands on their host, with infected sticklebacks having poorer body condition and lower energetic reserves [12,14], higher metabolic rate [15–17], and lower growth rate (in terms of lean mass) [12,14] than noninfected fish, with the strong deformation of the body expected to strongly constrain their mobility.

**Figure 1.**
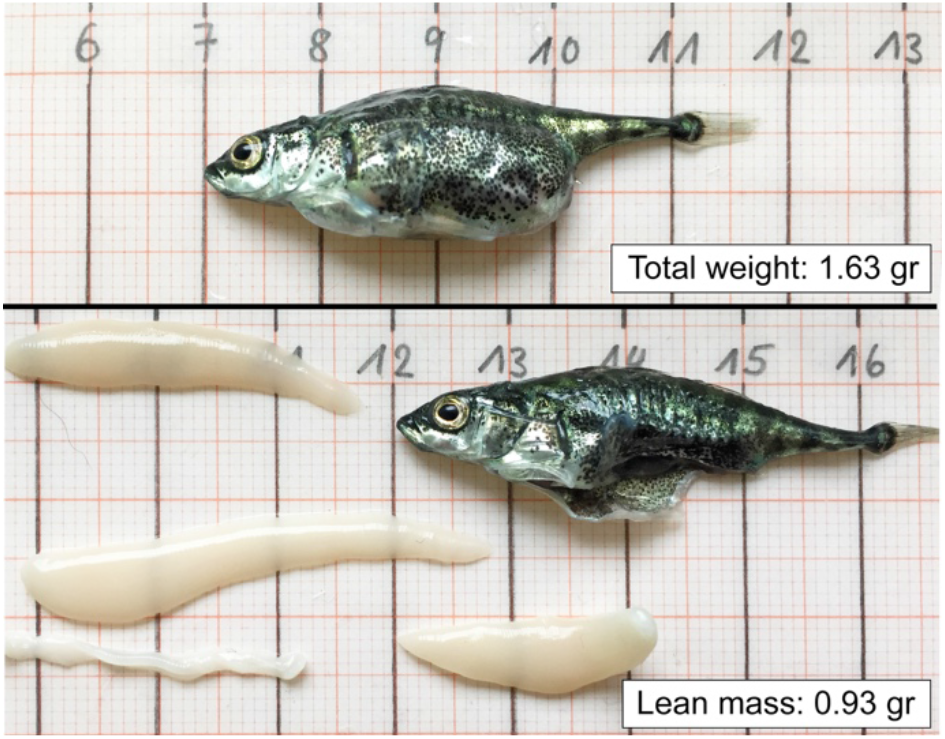
Photos of a euthanatized, heavily infected stickleback before (top) and after (bottom) the removal of the four *S. solidus* plerocercoids it carried.

Considerable work exists on the effects of *S. solidus* on the behaviour of infected sticklebacks, which reveals that the parasite strongly shifts the cost-benefit trade-offs of their host, with infected fish having reduced predator avoidance behaviour and increased motivation for food [18–27], with a higher parasite load resulting in stronger effects (e.g. [18,20,26]). In terms of mobility, researchers have long noted that infected sticklebacks move more slowly than noninfected fish (e.g. [10,14,16,18]), but the evidence is anecdotal. Furthermore, recent work investigating the total distance moved by freely moving sticklebacks did not find any effects [17,28]. Studies testing sticklebacks in forced swimming tests found that infected fish had reduced maximum swim speeds and endurance [16,29,30], and exhibit increasingly high metabolic costs when forced to move at greater speeds [15–17].

Overall, there is still little mechanistic understanding of the effects parasites like *S. solidus* have on the movements of their host [31]. Such knowledge is especially important because the behavioural changes that parasites may induce at the individual level may have large repercussions at the group level, with in turn wide-ranging ecological and evolutionary consequences [32]. Indeed, some studies, in fish in particular, have shown that parasites may affect within-group positioning and leadership, group cohesion and movement dynamics, and among-group assortment [6,7,33–39]. Furthermore, work on the stickleback - *S. solidus* system suggests that infected fish may have weaker social responses [33,37] that may influence the risk-taking behaviour of non-infected group mates [37]. But how exactly such social patterns emerge is not clear.

Here we systematically studied the fine-scale movements of individuals and pairs of sticklebacks with different levels of *S. solidus* infection to get a better understanding of how parasitism alters host movement dynamics and in turn influence social interaction patterns. We ran two experiments in which we tested fish individually and in pairs in an open environment and when being chased and startled, and ran individual-based simulations of self-organised groups to seek a parsimonious explanation of the observed collective patterns. We thereby specifically focused on parasite load (% of body weight) rather than just parasite status of the fish. We hypothesised that, as a result of lower movement capacity and increased energetic costs due to the parasite, infected fish would be slower and less mobile than non-infected fish and these effects to be more pronounced the higher the parasite load of the fish. We predicted that, due to self-organising effects, these individual-level changes would in turn result in pairs of infected fish to be slower, less cohesive and less coordinated than non-infected pairs, and mixed pairs to be primarily led by the likely faster, non-infected fish.

## Methods

### Subjects

Three-spined sticklebacks (*Gasterosteus aculeatus*) were collected from lake Constance, Germany using hand and dip nets, and subsequently housed in large social housing tanks (100 cm length x 50 cm width x 50 cm height), at the Limnological Institute, University of Konstanz. Tanks were connected to a flow-through system with water coming directly from the lake from a depth of 12 meters, resulting in a constant water temperature of ~ 13° C. Standard illumination was provided from above matching the day-night cycle. Fish were acclimated to the lab for at least three months before experiments. During this period, fish were fed defrosted bloodworms (Chironomidae) *ad libitum* once daily.

### Experiment 1

We randomly selected 180 fish from the social housing tanks, visually controlling for size (mean ± SE = 40.00 ± 0.32 mm), and moved them to our experimental lab. To be able to create specific pair combinations of infected and non-infected fish, individuals were anaesthetised using MS222, photographed from above using a tripod-mounted camera, and their infection status predicted based on their abdomen shape and size [11,13]. As no clear phenotypic differences exist between male and female sticklebacks under our laboratory conditions, the sex of the fish was not determined. To enable individual identification, fish were given a small coloured plastic tag with written id number on their first dorsal spine [40], after which they were left to recover in buckets of fresh, aerated water.

Fish were housed in five identical glass aquaria, each consisting of three compartments of equal size (20 × 40 × 40 cm; 34 cm depth), separated by mesh partitions, and with a fresh influx of lake water at 13 ± 1°C. Fish were randomly allocated to compartments in groups of 16 fish, and identifiable by their tag colour-id combination. After the experiments, all fish were euthanised with an overdose of MS222 (1g/l), weighed, their body cavity opened, and plerocercoids removed and weighed individually to the nearest mg. Based on the weight of the fish and their parasites we then determined the ‘parasite load’ for each fish as the percentage of body weight due to parasites [18,20,29] (for further descriptives, see Figure S1). Comparing the parasite status of the fish as determined pre- and post-mortem revealed a prediction accuracy of 97%, with all infected fish predicted correctly.

The experimental period started after fish were housed in the experimental holding tanks for five days, during which fish were fed defrosted bloodworms *ad libitum*, to allow for acclimation and stabilization of any social modulation effects [41]. Experiments were conducted using three identical white Perspex open-field arenas (50 × 70 × 10 cm; water depth: 8 cm). Arenas were placed inside a large, covered structure to minimise any external disturbances and provide diffuse lighting within. Trials were recorded from above with Raspberry Pi computers (RS Components Ltd) using pirecorder [42]. For experiment 1 we started by testing fish twice individually in the open arena (‘solo assay’; days 1 and 5). Fish were netted from their holding compartment and moved to the experimental arena where they were left to acclimate in a transparent cylinder (10 cm diameter) in the centre of the tank. After one minute, the trial started and the cylinder was remotely lifted by the experimenter, and fish were allowed to explore the arena for 10 minutes. After the trial fish were fed individually by moving them to a glass container containing two bloodworms before being returned to their housing compartment. The order of testing was randomised but the same order of testing was used for both testing days.

After two more rest days (day 9), fish were retested again but now in pairs (‘pair assay’). Individuals were randomly allocated to pairs based on their infection status to get roughly equal numbers of pairs in which no fish was infected (pair status: NN), only one fish was infected (pair status: MIX), and both fishes were infected (pair status: II). Due to some false negatives, determined from the post-mortem analyses (see above), and the loss of one fish, the actual pair numbers differed slightly between the treatment groups (n = 17, 21, and 21 pairs respectively). The proportion of individuals with low and high infected individuals was the same between the MIX and II pairs (45.1% and 45.9% respectively). As differences in body size have previously been shown to influence social interactions in fish [43], we further assorted fish based on their standard length (henceforth ‘body length’; BL), resulting in a negligible length difference between paired fish (1.66 ± 0.17 mm; see ESM Figure S2). Trials again lasted for 10 minutes.

### Experiment 2

To investigate how parasite infection alters the maximum movement capacity of individual fish (as outlined in [32]), we conducted an additional experiment and tested naïve fish under the demanding conditions of being chased and startled. 24 size-matched individuals (32.8 ± 0.4 mm), 11 of which were infected (confirmed post-mortem) were taken from the social housing tanks and moved to individual holding compartments (20 × 10 cm, enabling visual and chemical social cues) in our experimental lab. After two days of acclimation, each fish was subjected to a ‘startle assay’ and a ‘chase assay’ twice on two subsequent days (see ESM Figures S3 and S4). For the chase assay, fish were acclimated to the experimental tank for one minute after which they were briefly chased with a net moved at fishes’ maximum speed before being caught. For the startle assay we simulated a predator attack with a fake heron-like structure 50 cm above the water following previous work (e.g. [20,21,27]). Fish were released into the tank and after at least 30 seconds, the structure was remotely triggered the moment a fish entered that side of the tank, resulting in its immediate release into the water. Chase and startle tests were conducted under similar conditions as Experiment 1, i.e. an open, familiar Perspex tank (50 x 100 cm) in a covered structure, but trials were recorded with a GoPro camera fixed above the tank.

### Data processing

We used custom-developed tracking software to acquire detailed coordinate data of the fish as well as the orientation of their body during the trials. For the solo and pair assays, fish were automatically tracked while for the chase and startle assays this was done by manual tracking, resulting in four chase trajectories and two startle trajectories for each fish. Coordinates were converted from pixels to mm and subsequently smoothed using a Savitzky-Sgolay smoothing filter. After tracking, all trajectories were visually checked for any inconsistencies or errors.

From the tracking data, we computed each fish’s velocity, speed, acceleration, and turning speed. Measures linked to movement speed were computed in terms of body lengths to account for any effects of size. We estimated individuals’ movement tendency by computing the proportion of time spent moving, using a criterium of > 0.5 BL/s to determine if a fish was moving or not, based on the speed distribution, see Figure 2a. To estimate fishes’ cruising speed, we quantified their median speed when in this ‘moving state’. Neither the tendency to move or fishes’ movement speed were correlated with their distance from the tank walls (ESM Figure S5). As a measure of turning ability we computed the average time it took a fish to turn its body at least 15 degrees. To estimate movement predictability, we computed temporal autocorrelations of fishes’ speed and heading. For the chase and startle assay, we additionally computed the 0.75th quantile of fishes’ speed and acceleration (‘maximum’ speed/ acceleration).

**Figure 2.**
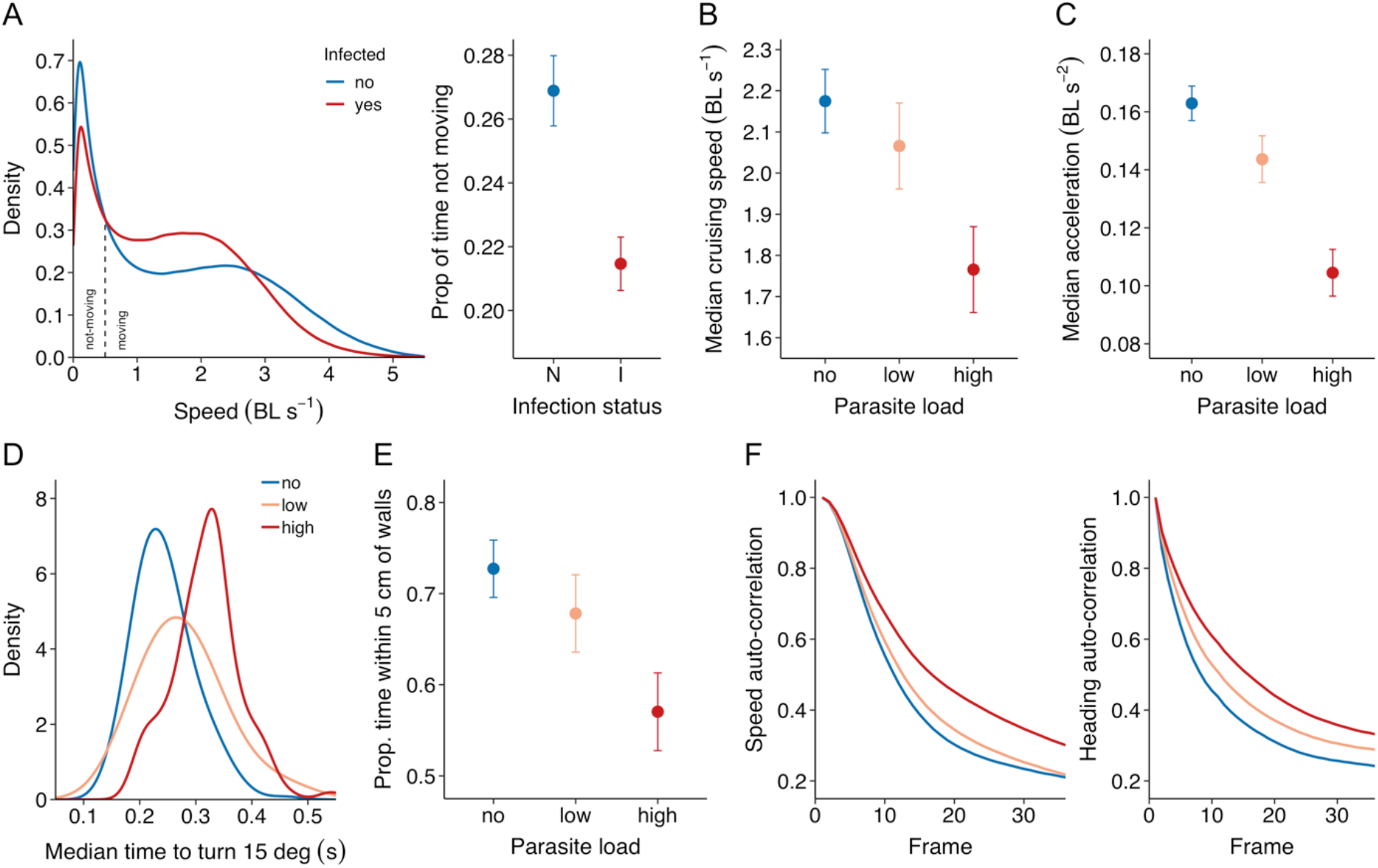
Movement characteristics of fish in the solo assay. A) Density plots of the speed distribution of infected (red; *n* = 94) and non-infected (blue; *n* = 86) fish (left), based on the full frame-by-frame dataset, and the proportion of time fish were in the not-moving state (< 0.5 BL s^-1^) in terms of their infection status (right). B) Fishes’ cruising speed, i.e. their median speed when in the moving state, and C) mean (tangential) acceleration for fish with no (*n* = 86), low (*n* = 47), and high (*n* = 47) parasite load. D) Density plots of the median time it took fish to turn their body at least 15 degrees, E) fishes’ ‘thigmotaxis’, the proportion of time spent within 5 cm of the nearest wall, in terms of their parasite load, and F) auto-correlation plots of fishes’ speed (left) and heading (right) in terms of their parasite load. Error bars are 95% confidence intervals of the mean.

For the pair trials we additionally computed the position and movement vector (between frames) of the group centroid. We then calculated each pairs’ cohesion, in terms of the distance between the two fish, alignment, in terms of the absolute difference in orientation, and group speed, in terms of the speed of the centroid. We also computed fishes’ front-back positioning relative to the group centroid, enabling us to determine which individual in the pair was more often in front. Finally, to determine the propagation of movement changes in the pairs, we ran temporal correlations on their speed and heading, and determined the average directional correlation between the two fish as a function of the delay in time [44,45]. A detailed explanation of the computation of these behavioural measures can be found in the ESM. For all variables we computed either the mean or median, based on the distribution of the raw data, and inter-quartile range on an individual- and pair-basis.

### Data analysis

To investigate the role of parasite infection on individual and pair movement dynamics we used a generalised linear mixed modelling (LMM) approach. The models of experiment 1 included as fixed effects trial (single assay only) and parasite load, categorised into a no *(n* = 84), low (0.002-0.27; *n* = 47), and high (0.27 - 0.49; *n* = 47) category (see ESM Figure S1 for further details). Fish id was added as a random factor, which was further nested within group id for the paired trials. For the chase and startle assays we ran separate models with 0.75th speed quantile and 0.75th acceleration quantile as response variables, infection status (noninfected, infected) and test (chase, startle) as fixed factors, and fish id nested within trajectory as a random factor. In general, we used models fitted with a Gaussian error distribution except for count data, which was fitted to a Poisson error distribution with log-link function. Minimal adequate models were obtained using backward stepwise elimination, i.e., sequentially dropping the least significant terms from the full model, until all terms in the model were significant. Statistics for non-significant terms were obtained by adding the term to the minimal model. Neither housing compartment nor tank number had an effect on any of the behaviours analysed. Residuals were visually inspected to ensure homogeneity of variance, normality of error and linearity where appropriate; data was square-root or log-transformed otherwise. Means are quoted ± SE unless stated otherwise. All data were analysed in R 3.5.0.

### Individual-based models

We engaged individual-based simulations of self-organised groups to further investigate the underlying mechanisms of how parasitism likely drives group-level effects and to seek a parsimonious explanation for the observed patterns of cohesion, alignment, and leadership. Motivated by the behavioural differences observed in the experimental data, we simulated two individuals and integrated heterogeneity in speed, turning ability, and stop-go tendency. A mathematical description and details of the model can be found in the ESM.

## Results

### How does parasite infection affect individual movement?

In the solo assay, infected fish were less variable in their movement speed than non-infected fish and both lower maximum speeds and a higher tendency to keep moving (Figure 2A). Parasite load had a strong effect on both the cruising speed (median speed when moving; χ^2^ = 35.68, *p* < 0.001) and mean acceleration (χ^2^ = 100.3, *p* < 0.001) of the fish, with heavily infected fish swimming and accelerating more slowly than slightly- and non-infected fish (Figure 2B, 2C, ESM Figure S6). Fishes’ body weight did not affect acceleration (χ^2^ =1.20, *p* = 0.273).

Although parasite load did not influence the maximum turning rate of the fish on a second-by second basis (χ^2^ = 1.33, *p* = 0.514), heavily infected fish took longer to make a more abrupt change in their orientation (15°) than slightly- and non-infected fish (χ^2^ = 52.60, *p* < 0.001; Figure 2D). Parasite load also affected fishes’ thigmotaxis, with non-infected fish spending the most and highly infected fish the least amount of time within 5 cm of the tank walls (χ^2^ = 31.59, *p* < 0.001; Figure 2E).

Parasite load had a strong negative effect on how quickly fishes’ movements changed over time, both in terms of their speed (χ^2^ = 53.01, *p* < 0.001) and heading (χ^2^ = 81.51, *p* < 0.001): non-infected fish were the least predictable and highly infected fish the most predictable in their movements (Figure 2F). The effect of predictability in turning behaviour remained when looking at how quickly fish changed their heading over distance rather than over time, which is affected by speed, although the effect was less strong (χ^2^ = 13.39, *p* = 0.001).

In the chase and startle assays, fish exhibited on average much higher speeds and acceleration compared to in the solo assay (Figure 2, 3). However, also under these high-demand conditions, infected fish were found to be the slowest, reaching a significantly lower maximum speed (χ^2^ = 24.33, *p* < 0.001) and lower maximum acceleration (χ^2^ = 29.14, *p* < 0.001) than non-infected fish (Figure 3, ESM S3, S4). Non-infected fish also reached their maximum speed much earlier than non-infected fish when being chased and were able to swim considerably further away in the first moments after being startled (ESM Figure S7).

**Figure 3.**
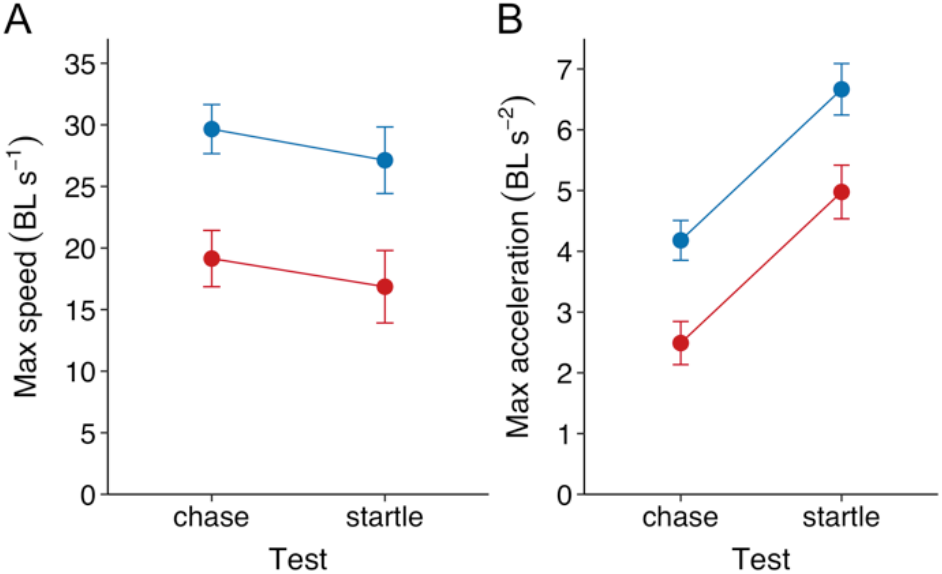
A) Maximum speed and B) maximum acceleration in terms of fishes’ body length for infected (red; *n* = 11) and non-infected (blue; *n* = 13) fish in the chase and startle assay. Maxima were quantified based on .75 quantile probabilities and error bars are 95% confidence intervals of the mean.

### How does parasite infection affect social interactions?

Although fish strongly conformed in their speed, relative inter-individual differences in speed were maintained (*R*_C_ = 0.62, 95% CI: 0.49-0.72; see ESM Figure S8). The movement speed of the pairs was significantly affected by pairs’ infection status (*F*_2,56_ = 15.59, *p* < 0.001), with those consisting of two infected fish swimming significantly slower than pairs in which only one fish was infected, which in turn were slower than pairs of non-infected fish (Figure 4A). Despite considerable variation in size among the pairs (see ESM Figure S2), body size did not play a significant role (ΔAIC = 1.43). As in the solo assay, infected fish swam more slowly than non-infected fish, but only so in the homogeneous pairs (status × relative partner status: χ^2^ = 17.95, *p* < 0.001), showing that both infected and non-infected fish in the mixed pairs adjusted their swimming speed in response to their partner (Figure 4B). Such an effect was not found for fishes’ acceleration (χ^2^ = 0.63, *p* = 0.427) and instead, infected fish acceleration was lower than that of non-infected fish irrespective of their partner (χ^2^ = 56.92, *p* < 0.001; Figure 4B). In general, fish exhibited different social response patterns depending on their infection status (ESM Figure S9), with infected fish showing smaller speed and turning changes in response to their partner.

**Figure 4.**
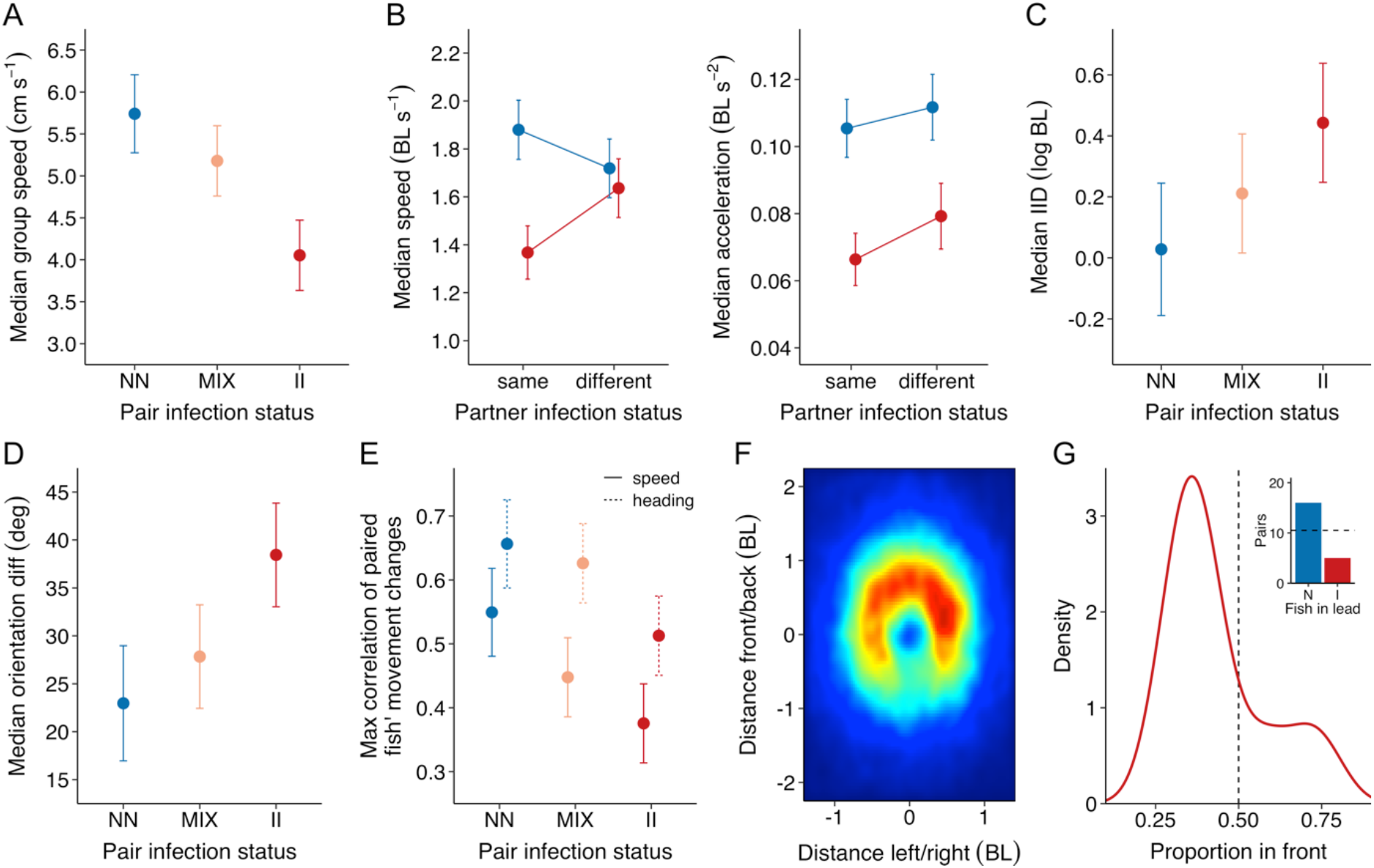
Movement characteristics of fish in the pair assay. A) Median movement speed of the group centroid in cm/s for pairs in which neither fish was infected (NN, *n* = 17), one fish was infected (MIX, *n* = 21) of both fish were infected (II; *n* = 21). B) Fishes’ median cruising speed in BL/s and median acceleration in BL/s^2^ in terms of fishes’ infection status and the relative infection status of their partner. C) Relationship between pairs’ infection status and their median inter-individual distance in average body length (log-transformed), D) their median difference in orientation, and E) the max correlation coefficient of fishes’ speed (solid line) and heading changes (dashed line). Error bars are 95% confidence intervals. F) Heatmap of the spatial positioning of infected fishes’ partner in the mixed pairs based on the frame-by-frame data. Data is cropped to 73% of the full parameter space to show the most relevant area only, with the colour scale being proportional to the densest bin of the plot (blue = low; red = high). G) Density plot of the proportion of time infected fish spent in front in the mixed pairs, with inset showing the number of pairs for which the non-infected (N) or the infected (I) fish predominantly led (*n* = 21 mixed pairs).

The infection status of the pair affected their cohesion (*F*_2,56_ = 4.13, *p* = 0.021, Figure 4C) and alignment (*F*_2,56_ = 7.97, *p* < 0.001, Figure 4D), with pairs of non-infected fish being the most cohesive and aligned and pairs of two infected fish the least so. Furthermore, while at very low speeds non-infected pairs still stayed relatively well aligned, this was less the case for infected pairs (ESM Figure S10). Pairs’ infection status also significantly affected their coordination, both in terms of how paired individuals modified their speed in response to their partner (*F*_2,56_ = 7.09, *p* = 0.002) and in terms of their orientation (*F*_2,56_ = 5.62, *p* = 0.006), with infected pairs being the least and non-infected pairs the most coordinated (Figures 4E). Spatial leadership was affected by infection status, with mixed pairs being predominantly led by the, generally faster, non-infected fish (Figure 4F, 4G), both in terms of the number of pairs (χ^2^ = 5.76, *p* = 0.016) and the average proportion of time spent in front (*t*_20_ = 1.93, *p* = 0.034). This was strongly linked to differences in speed (ESM Figure S11), with infected fish swimming more slowly when in the front position compared to non-infected fish (ESM Figure S10A).

### Individual-based simulations

The simulations (see ESM for detailed results) showed that both high values of turning responsiveness and low values of turning noise considerably increased pair alignment. A similar but weaker effect was observed in terms of cohesion. Spatial leadership was positively correlated with individual speed and negatively correlated with turning responsiveness and proportion of time in the stop state. But while all three factors influenced leadership, speed had the strongest effect and could offset opposite effects arising from the other two factors.

## Discussion

Using the stickleback - *S. solidus* model system, this study aimed to gain a mechanistic understanding of how parasite infection affects individual movement ability and thereby shapes group-level patterns. When tested individually, infected fish moved and accelerated more slowly than non-infected fish, both when free to move and when chased or startled. Infected fish also moved more continuously and were more predictable in their movement changes than non-infected fish. Although we used naturally infected fish, we found that all effects were strongly positively linked with parasite load, with larger impairments in behaviour the higher fishes’ parasite load. In turn, parasite infection strongly affected fishes’ social interactions, with infected pairs moving more slowly and being less cohesive, less aligned, and less coordinated than pairs of non-infected fish, and mixed pairs being predominantly led by the non-infected individual. Individual-based simulations of pairs demonstrate these social patterns can emerge from individual differences in speed, turning ability, and stop-go behaviour that arise due to parasite infection, with speed being the predominant factor.

The median movement speed of infected and non-infected fish was very similar across the solo trials, in line with previous work [17,28]. This may lead to the conclusion that parasite infection does not affect individual movement. However, detailed movement analyses revealed that speed distributions were bimodal, with two peaks corresponding to a ‘stop’ and ‘moving’ states. In addition to spending more time in the moving state, infected fish also moved at significantly lower cruising speeds when compared to non-infected fish and did not accelerate as strongly. These differences may be explained by the combination of energetic and physical constraints imposed by the parasite. First of all, *S. solidus* increases its host’s metabolism [15–17] and as a result fish may voluntarily move at lower speeds to save energy. Secondly, the parasite grossly distends the body of their host, which impairs its streamlining and increases drag [12,16]. In line with the mechanical law that drag increases with velocity squared [46], infected fish have previously been shown to have much higher increases in metabolism with increasing speed compared to non-infected fish [16,17], explaining why infected fish may not be able to sustain higher speeds. Third, *S. solidus* increases its host’s body volume and specific weight, shown here (see Supplementary Material) and in previous work [14–16]. As the forward thrust required to change the inertia of a body in water is proportional to its volume and weight [47], infected fish are likely physically constrained in terms of their acceleration, and such changes would be energetically more costly [46]. Furthermore, the distended abdomen of infected fish also increases its rigidity and thus may thereby restrict propulsive movement [10] by impairing their ability to flex the anterior and posterior sections of their bodies [48]. These mechanisms may also help explain why infected fish were more consistent in their speeds and less likely to stop moving completely (see also [23]), and increasingly so with higher parasite load.

When chased or startled, infected fish exhibited considerably lower maximum swimming speeds and acceleration than non-infected fish (see also [29,30]). This again highlights the physical constraints imposed by the parasite that can negatively influence the performance capacity of individuals (see [32]). Indeed, experimental work with artificial parasites has revealed that it is especially the drag effects rather than metabolic effects that are important for reduced responses under such highly demanding conditions [49]. The finding that infected sticklebacks fled shorter distances after a simulated predator attack (see also e.g. [20]), may thus not necessarily (only) imply reduced risk-taking behaviour but could also be the result of a mechanical effect of the parasite. Besides affecting the speed, acceleration, and movement consistency of the fish, parasite infection was also found to affect turning behaviour, with infected fish showing slower turning, with again the effect being more pronounced for more heavily infected fish. Body rigidity is known to be detrimental to manoeuvrability and turning performance [47,50]. Therefore, our results are likely to be the result of the increased rigidity of the grossly distended body of infected fish. This effect may potentially also help explain the observed finding that infected fish had weaker thigmotaxis (see also [21]), since increased rigidity would compromise fishes’ ability to turn in response to a nearby wall.

Parasite-induced effects on individual movement were found in turn to influence social movement patterns. Besides pairs of infected fish tending to be slower than pairs of noninfected fish, they were also less cohesive, in line with previous work on fish infected with parasitic worms [7,33,34], and less aligned, with mixed pairs showing intermediate levels of these behaviours. Such patterns can emerge naturally from the impaired mobility of infected fish (lower speed/acceleration/manoeuvrability, see above) by the constraints it places on the individual to conform with its partner. This is supported by our analyses of the social response patterns, which showed that infected fish had weaker/slower responses to their partner than non-infected fish, and our simulations, which revealed that a lower turning responsiveness decreased group cohesion and alignment. Although the simulations do not differentiate between the physiological ability and social motivation to turn, the observation that infected fish were slower in turning in the solo trials suggests that this difference in responsiveness is likely more biomechanical compared to a lower social motivation. This is in line with the finding that pairs of non-infected fish not only had higher alignment when moving, which could arise directly from their higher speed [45,51], but also when moving at very low speeds. Another potential explanation for the observed group-level patterns is that, because the parasite increases their host’s metabolism [17], infected sticklebacks may have a higher motivation to search for food, since sticklebacks searching for food tend to be less cohesive [52]. However, in that case also their individual and group speeds would be expected to be higher, and fish never experienced food in the testing environment. The abdominal swelling itself may also directly impair the ability of infected fish to accurately assess proximity (see also [34]), and explain why infected fish respond more slowly to, and stay further away from, others and the boundaries of their environment. It is important to note that parasites like *S. solidus*, which is not directly transmittable, will have very different effects on the trade-offs of grouping of their hosts than directly transmittable parasites, and future experiments are needed that compare different host-parasite systems.

Parasite infection had a strong effect on social coordination, with infected pairs being the least, and non-infected pairs the most coordinated. While the lower coordination of infected pairs may be a result of a reduced sociability of infected fish, the results of the solo and pair context together strongly suggest infected fish have a physical impairment that negatively impacts their capacity to respond to others’ movements and thereby reduces coordination. The consideration of such parasite effects is important as they may strongly affect the transfer of information and may thus influence group functioning [53]. The finding that mixed pairs were primarily led by the non-infected fish is explainable by self-organising effects from individual differences in speed induced by the parasite. Indeed, both our empirical results and our simulations showed a positive relationship between leadership and individual speed, conform previous work (e.g. [45,54,55]). The simulations also revealed that a lower turning responsiveness and a higher tendency to keep moving, aspects that characterise infected fish, both actually increased leadership behaviour [54]. While the present study was purposefully conducted on the smallest group of two individuals, our results provide clear predictions for the effects of *S. solidus* on larger groups and among-group dynamics, with infected fish predicted to be less able to form stable, cohesive, coordinated groups, more likely to occupy positions on the edge and back of groups, and more likely to be isolated (see also [54]).

Many studies have previously shown that sticklebacks infected by *S. solidus* exhibited reduced predator avoidance behaviour and take more risks than healthy individuals [19,20,22,26,27], which may enhance the transmission of the parasite to its final host. Here we provide quantitative results of two additional, rarely considered factors that may increase the predation risk of infected fish (see e.g. [56]). Specifically, we show that infected fish are less likely to stop moving completely and are slower to change in their speed and heading over time, as revealed by autocorrelation analyses, and are therefore more predictable in their movements. As a result, predators may both more easily detect infected compared to healthy individuals and better able to predict their future position. This may be especially important for many piscivorous birds that hunt from above, the final host of *S. solidus*. Together, our findings thus suggest that the changes in host morphology and physiology caused by the parasite lead to changes in individual and social behaviour that may in turn increase chances of transfer to its final host. Although our findings do not provide direct evidence for or against direct manipulation, they suggest such more complex mechanisms may not necessarily need to evolve as the parasite can exploit the cascading effects caused by its changes on host physiology and biomechanics. This is supported by our finding that the observed effects of the parasite increase with increasing parasite load, in line with previous work [31], rather than a clear switch point in weight linked to infectivity [57], which would be expected under such direct parasite manipulation effects. Further empirical and theoretical work is needed that properly considers both active and passive mechanisms for parasite effects on host behaviour across different host-parasite systems. A potential shortcoming from our study is that we worked with naturally infected sticklebacks as it might be hypothesised that different phenotypes might also differ in their likelihood to be infected. We focused our analyses primarily on parasite load, which is not likely to be strongly influenced by intrinsic phenotypic differences. Furthermore, some data exists that suggests at least body condition parameters do not differ between sham and exposed, but not infected fish [58]. Future work in which fish with different behavioural phenotypes and immune status are experimentally infected to reveal if such differences may interact with the susceptibility and subsequent effects of parasite infection [9].

This year it is exactly 100 years since the complete complex life cycle of *S. solidus* was discovered [59]. While during this period we have gained many important insights into this fascinating host-parasite model system, much about the effects of the parasite on the individual and social movements of its host has remained unclear. By systematically studying the movement dynamics of differently infected sticklebacks using a combination of empirical and theoretical approaches, we provide new insights into the effects of *S. solidus* on the movements and social behavioural patterns of its host. These findings also contribute to the general understanding of the mechanisms by which parasites may affect host movement dynamics and the way in which individual differences in movement capacity can shape and impair group-level patterns. Exciting future work lies ahead to further investigate how parasitism may affect organismal movement at different social scales and thereby potentially shapes the structure and assortment of animal groups.

## Acknowledgements

We thank Jörn Scharsack and Nicolle Demandt for helpful feedback, Jonas Bleilevens for help with experiments, and Myriam Schmid for assistance with fish holding

## Ethics

This research was performed following the guidelines for the Use of Animals in Research of the Association for the Study of Animal Behaviour (ASAB/ABS). Animal care and experimental procesures were approved by Regierungspräsidium Freiburg under animal ethics permits G-16/144 and G-19/50.

## Data accessibility

Data accompanying this paper can be found in the Mendeley Data repository (doi: 10.17632/gjr8gxk6nw.1). Python code for running the individual-based simulations can be found on GitHub (https://git.io/Je6OT).

## Author contributions

The study was conceived by J.W.J. and G.P.F.M. with helpful input from all other authors. G.P.F.M. and J.W.J. performed the experiments with support from J.B.-G., J.D and J.W.J developed and performed the model simulations, J.W.J. processed and analysed the data, J.W.J. drafted the manuscript with substantial feedback from all other authors.

