## Supplementary Material for "*Schistocephalus* parasite infection alters sticklebacks’ movement ability and thereby shapes social interactions"

### ELECTRONIC SUPPLEMENTARY MATERIAL

#### Appendix A: Supplementary Experimental Methods

##### Data processing and analysis

*Computation of individual characteristics*

From the discrete tracking data, we determined each fish's velocity, speed, turning speed, and relative position to the group centroid over time using the following series of calculations. With the vector $\mathbf{r}_i(t) = (x_i(t), y_i(t))$ denoting the position of fish $i$ at time $t$, we approximated its velocity vector $\mathbf{v}_i(t)$ using the forward finite difference:

$$\mathbf{v}_i(t) = \frac{\mathbf{r}_i(t + \Delta t) - \mathbf{r}_i(t)}{\Delta t},$$

where $\Delta t$ = 1/24 s is the time interval between subsequent position measurements. A fish's speed $\mathrm{s}_i(t)$ was then approximated by the norm of the velocity vector:

$$s_i(t) = |\mathbf{v}_i(t)|.$$

Following from this calculation, we determined the change in a fish's speed over time as:

$$\mathrm{a}_i(t) = \frac{s_i(t + \Delta t) - s_i\mathrm{v}_i(t)}{\Delta t}.$$

Next, we estimated a fish's angular velocity (i.e. turning speed) at time $t$ based on changes in the orientation (heading) angle $\theta_i$ rather than the direction based on its velocity vector, which becomes spurious at low speeds. The angular velocity is thus calculated from the data as

$$\omega_i(t) = \frac{\theta_i(t + \Delta t) - \theta_i(t)}{\Delta t}$$

and then corrected to ensure that values fall within the range $-\pi < \omega_i \leq \pi$. Subsequently, we determined the group centroid, i.e. mean coordinates of both fish in a group, as $\mathbf{r}_c(t) = (x_c(t), y_c(t))$ and calculated the distance of each fish to the group centre

$$CD_i(t) = \sqrt{(x_i(t) - x_c(t))^2 + (y_i(t) - y_c(t))^2}.$$

To further estimate fish' turning ability, for each frame in the time series we moved forward in time until the relative absolute angle difference between those two frames was more than a threshold value, recorded the latency in terms of frame number, and then computed the median turning latency, using a threshold of 15 degrees.

*Computation of spatial positioning*

To determine the front-back positioning of the pair of fish, we shifted their $(x, y)$ coordinates such that the origin of the coordinate system was at the group centroid. We then determined the angle between the positive $y$ axis through the group centroid and an individual's position as

$$\delta_i(t) = atan2\big(x_i(t) - x_c(t), y_i(t) - y_c(t)\big),$$

which we used to calculate an individual's relative direction to that of the group centre

$$\sigma_i(t) = \delta_i(t) - \psi_c(t),$$

where $\psi_c(t)$ is the heading angle of the group centroid. Based on the relative direction and distance to the group centre, we then calculated the relative position for each fish in a pair to the centroid

$$(x_i', y_i') = CD_i(t)\big(\sin(\sigma_i(t)), \cos(\sigma_i(t))\big).$$

These transformed coordinates enabled us to determine spatial leadership positioning of the fish, with a positive $y'$ indicating a fish was in front and a negative $y'$ indicating a fish was behind the group centroid. We then counted the proportion of frames that each fish was located in the forward-most position of the pair.

*Computation of group-level measures*

As group-level measures we computed group cohesion as the distance between the two fish:

$$c_{ij}(t) = \sqrt{\big(x_i(t) - x_j(t)\big)^2 + \big(y_i(t) - y_j(t)\big)^2},$$

computed alignment in terms of fishes' absolute difference in heading via:

$$\varphi_{ij}(t) = |\theta_{i-}\theta_j|$$

and

$$\{\varphi_{ij}(t) > \pi\}: \varphi_{ij}(t) = 2\pi - \varphi_{ij}(t),$$

and defined group speed $s_c(t)$ by the norm of the centroid velocity vector:

$$s_c(t) = |\mathbf{v}_c(t)|$$

*Temporal correlations*

To investigate predictability in fishes' movements, we examined temporal auto correlations in terms of the speed $s_i$ and heading $\theta_i$ of the fish. To do so, we created two copies of the time series data subsetted the second time series up to a delay $d$ of 96 frames. We determined the correlation for each delay $C_i(d)$ and computed the auto-correlation coefficient $\tau$ based on the slope of the resulting function. Fish with a greater auto-correlation change less quickly in their behaviour over time and are thus more predictable than those with a smaller $\tau$.

To investigate leadership in terms of how individuals guided the motion of their partner, we used a similar approach, but now instead of comparing the data of a fish against itself, we compared it to the data of its partner, increasingly delayed in time (see also [1–3]). We created two sets of time series of each fish's speed and heading, subsetted the dataset of the other fish again up to a delay $d$ of 96 frames, and determined the correlation for that time step $C_{ij}(d)$. Over all timesteps, we then identified the maximum value of $C_{ij}(d)$ and delay $d$ that corresponds to this maximum. A positive value suggests that fish $j$ adjusted its motion to match that adopted by fish $i$ at an earlier time, or, in other words, that fish $j$ was copying the speed/orientation of fish $i$.

### Appendix B: Supplementary Experimental Results

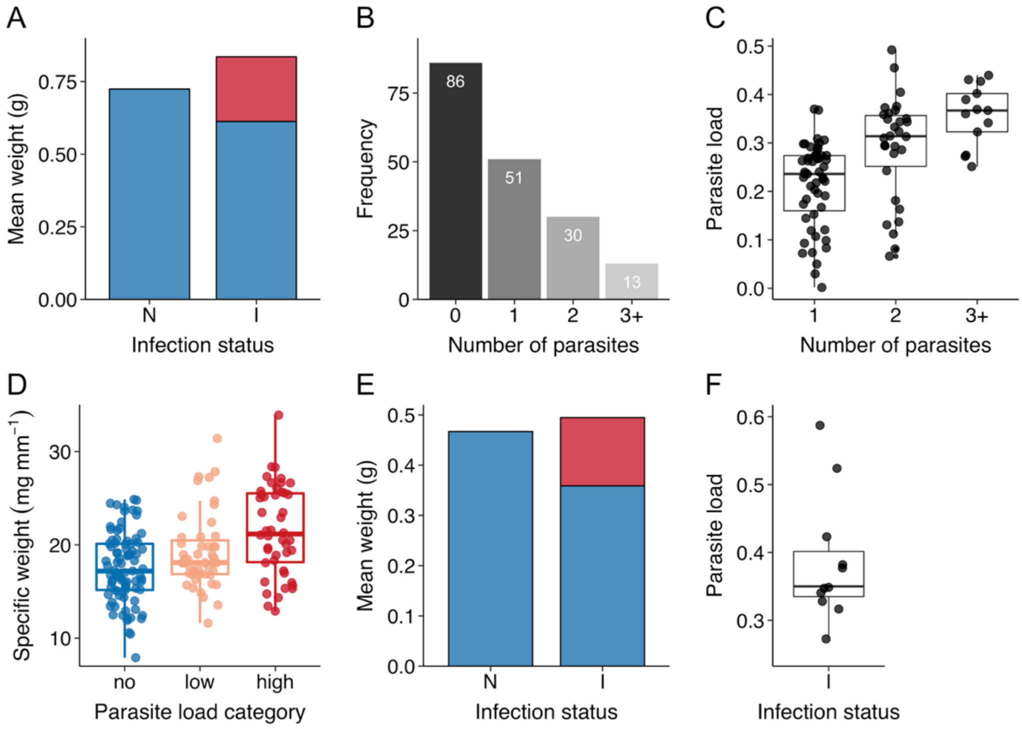


**Figure S1.** Detailed characteristics of the fish used in the first (A-D) and second (E-F) experiment. A) Mean weight of non-infected (N; $n$ = 86) and infected (I; $n$ = 94) fish in terms of fishes' own body mass (blue) and the mass of their parasites (red). B) Frequency table of the number of parasites per fish with in white the fish count. Fish had an average of 1.69 ± 0.10 parasites that ranged up to a maximum of 6. C) Relationship between the number of parasites and fishes' parasite load, i.e. the proportion of body weight attributable to the parasite(s). D) Relationship between the parasite load and fishes' specific weight for the no ($n$ = 86), low ($n$ = 47), and high ($n$ = 47) category. Although there was no difference in standard length between infected and non-infected fish ($t_{176.4}$ = -1.60, $p$ = 0.111), infected fish were significantly heavier in terms of total body weight than non-infected fish ($U$ = 4794, $p$ = 0.021), but lighter in terms of body mass (i.e. excluding the parasites; $U$ = 2764, $p < 0.001$). As a result of the extra weight of the parasite(s), infected fish had a higher relative mass in terms of body size compared to non-infected fish (infection status × total weight interaction: $F_{3,173}$ = 12.71, $p < 0.001$). While body size was positively correlated with absolute parasite load ($r_s$ = 0.52, $p < 0.001$), parasite load in terms of the proportion of weight of the fish, which ranged from 0.002 to 0.492, was not affected by size ($r_s$ = 0.03, $p$ =0.805). E) Mean weight of non-infected (N; $n$ = 13) and infected (I; $n$ = 11) fish used in experiment 2 in terms of their own body mass (blue) and that of their parasites (red). F) Boxplot showing the parasite load of the infected fish. Plots A and E, and C and F respectively show that the fish used in the two experiments are very comparable, and although a bit smaller and lighter in the second experiment, those fish also had a slightly higher total body mass but lower mass excluding the parasite.

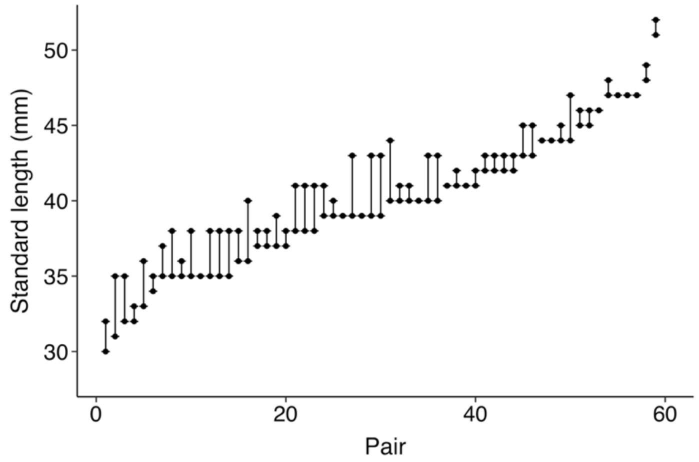


**Figure S2.** Figure showing the size-assortment for the fish into experimental pairs ($n$ = 59 pairs).

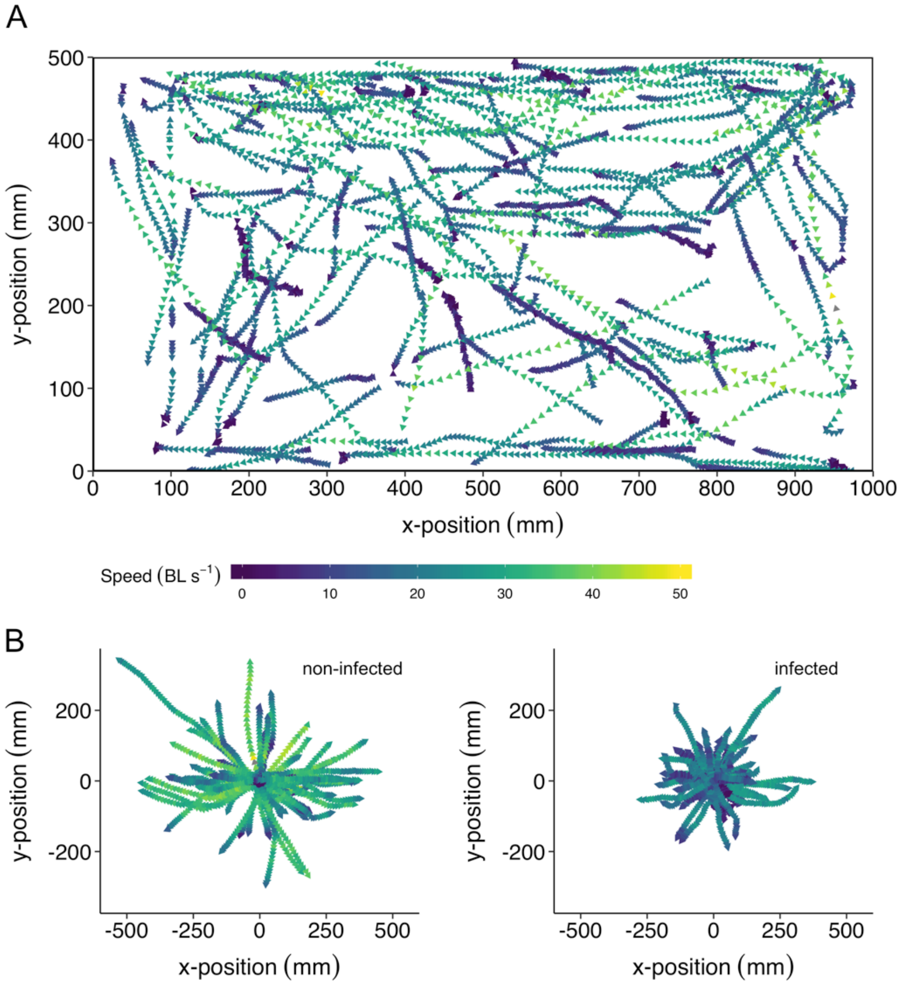


**Figure S3.** Visualisations of fishes' trajectories in the chase assay, with speed coloured from dark blue (slow) to yellow (fast). A) Trajectories with absolute coordinates in the tank. B) Trajectories of non-infected (left; *n* = 13 fish) and infected (right; *n* = 11 fish) fish (72 and 59 trajectories respectively) with relative coordinates to the starting position of each individual chase.

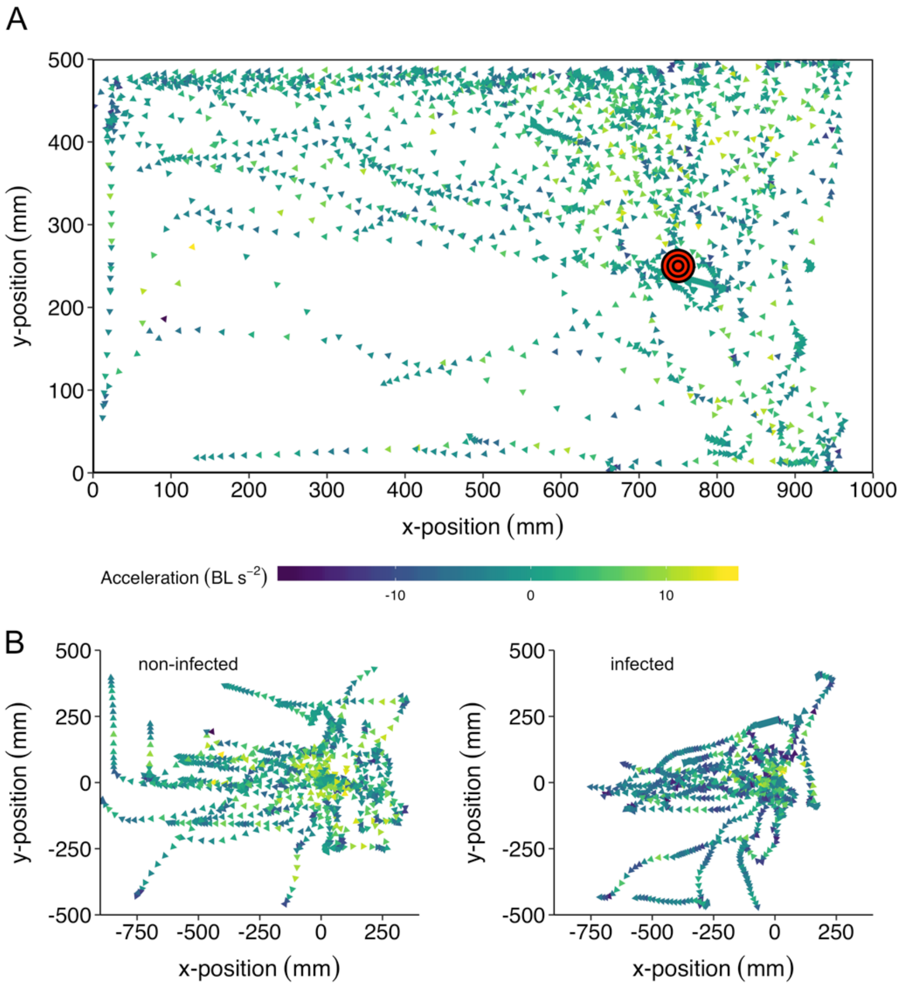


**Figure S4.** Visualisations of fishes' trajectories in the startle assay, with acceleration coloured from dark blue (strong deacceleration) to yellow (strong positive acceleration). A) Trajectories with absolute coordinates in the tank with the point of impact indicated by the red and black circle. B) Trajectories of non-infected (left; *n* = 13 fish) and infected (right; *n* = 11 fish) fish (25 and 22 trajectories respectively) with relative coordinates to their starting position when the fake heron head was released.

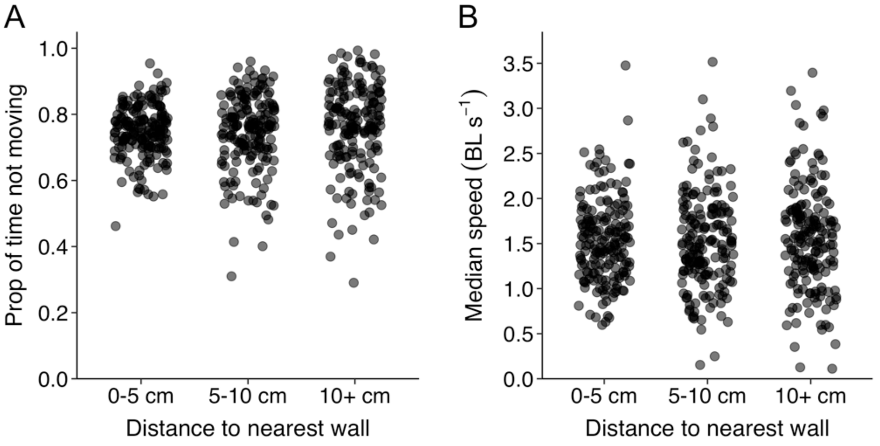


**Figure S5.** Scatterplots of A) the proportion of time fish (*n* = 179) were moving (> 0.5 BL/s) and B) fishes' median movement speed (in BL/s), with the data of experiment 1 subsetted to fish being within 5 cm of the nearest wall, 5-10 cm from the nearest wall, or more than 10 cm from the nearest wall (maximum possible distance is 25 cm).

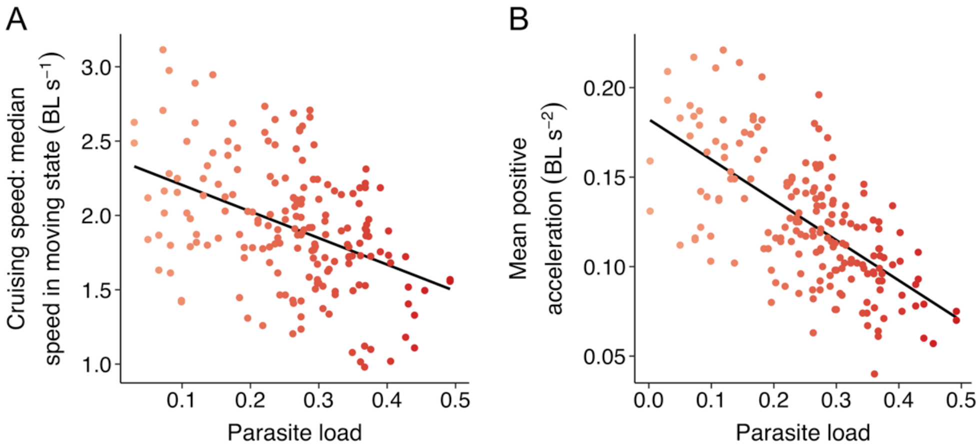


**Figure S6.** Relationship between parasite load and A) fishes' median speed when in the moving state (i.e. moving at > 0.5 BL/s), and B) fishes' mean positive (tangential) acceleration during the two solo trials (*n* = 94 fish).

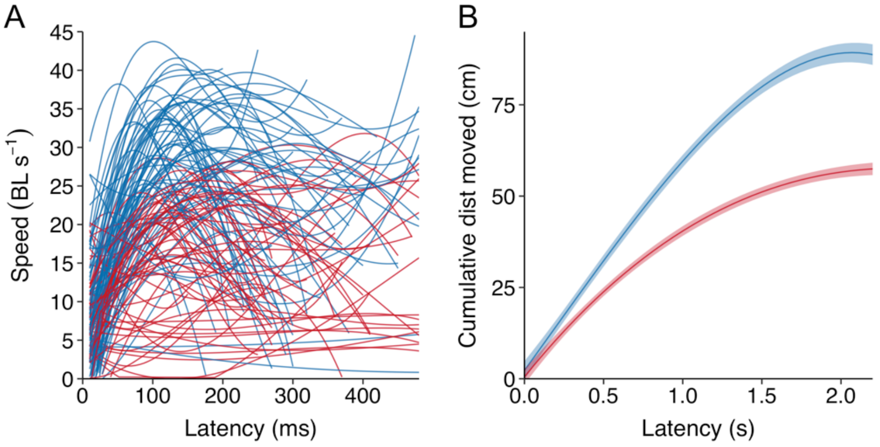


**Figure S7.** A) Smoothed polynomials of fishes' speed over time for individual trajectories during the chase assay, with infected fish in red (*n* = 72 trajectories of 13 fish) and non-infected fish in blue (*n* = 59 trajectories of 11 fish). B) Smoothed polynomials of the cumulative distance moved for infected (red) and non-infected (blue) fish in the startle test from the time the fake heron head impacted the water.

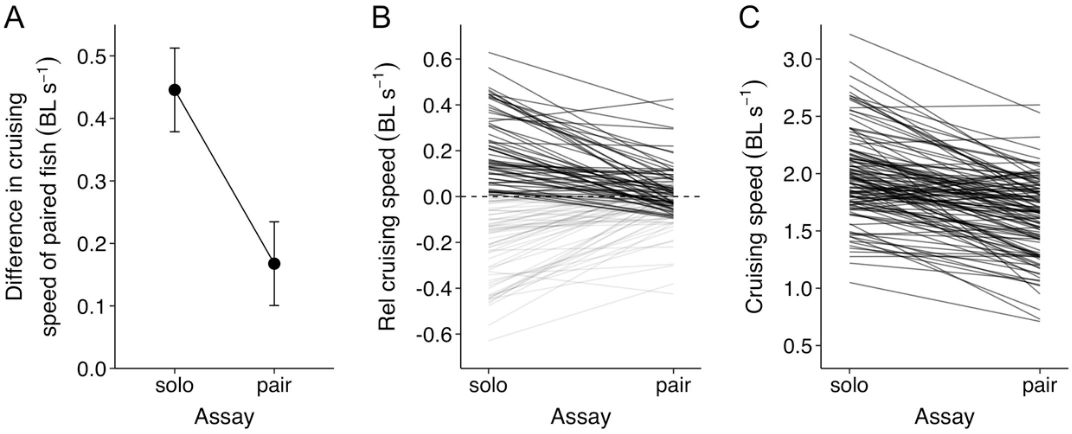


**Figure S8.** A) Difference in cruising speed between the two paired fish in the solo and pair assay. B) Same data as for A but now with the speed of the individual fish plotted relative to that of their partner. As fishes' relative speed is opposite to that of their partner, negative relative cruising speeds are semi-transparent. This plot also shows that fish that were faster in the solo assay also generally remained the faster fish in the pair assay. C) Fishes' absolute cruising speed (speed when moving at least 0.5 BL/s) in the solo and pair assay, showing high inter-individual differences, as indicated by the parallel lines. For all plots *n* = 118 fish and 59 groups. Data of the solo assay is based on the mean across the two trials for each fish.

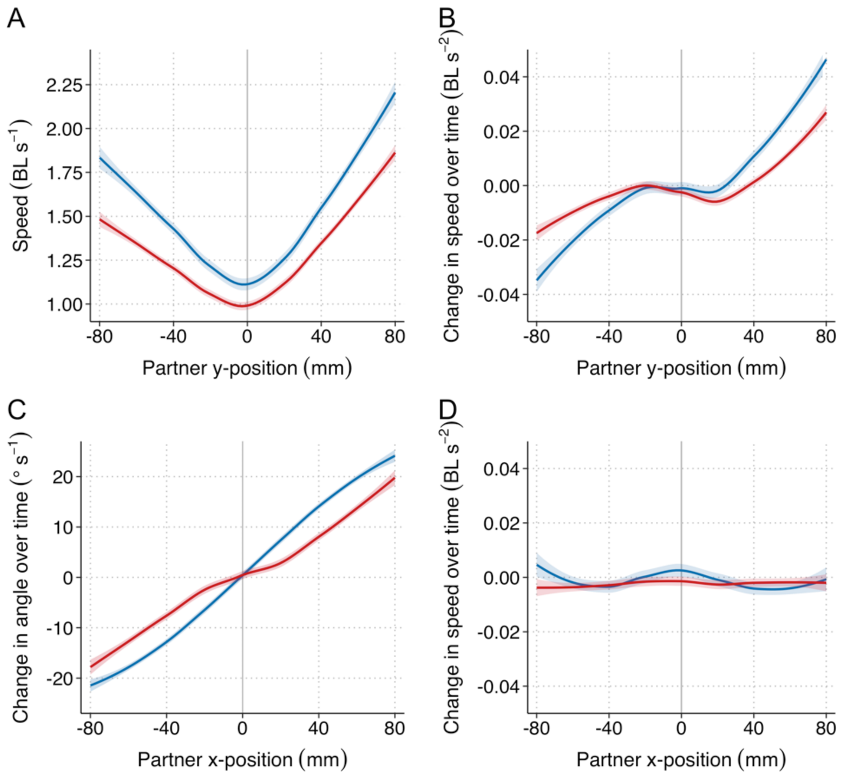


**Figure S9.** A) The mean speed, B) mean change in speed over time, C) mean change in orientation angle over time, and D) mean change in speed over time when both fish were approximately side by side, for infected (red) and non-infected (blue) fish as a function of the relative y-position (front/back) of their partner (A & B) or left-right position (C & D). Plots B and C describe how individual fish adjust their velocity as a function of the relative position of their partner. Ribbons indicate 95% confidence intervals. The data of the pair trials was subsetted to instances where both fish were within 100 mm from one another (80.4% of the full data) and for plot D further subsetted to instances where fish were less than 20 mm in front of one another (31.4% of the full data). These plots show that while non-infected fish were generally faster than infected fish, non-infected fish tended to adopt lower speeds when their partner was behind them (A). Furthermore, both fishes adjusted their speed depending on their partner's position, however, infected fish responded less strongly than non-infected fish (B). While both fish showed a strong tendency to turn towards their partner, the magnitude of turning changes was less strong for infected fish, although the effect was weaker than that in terms of speeding changes (C). Finally, when fish were approximately side-by-side, non-infected fish tended to accelerate and thereby take the front-most position, whereas infected fish tended to continue at their current speed (D).

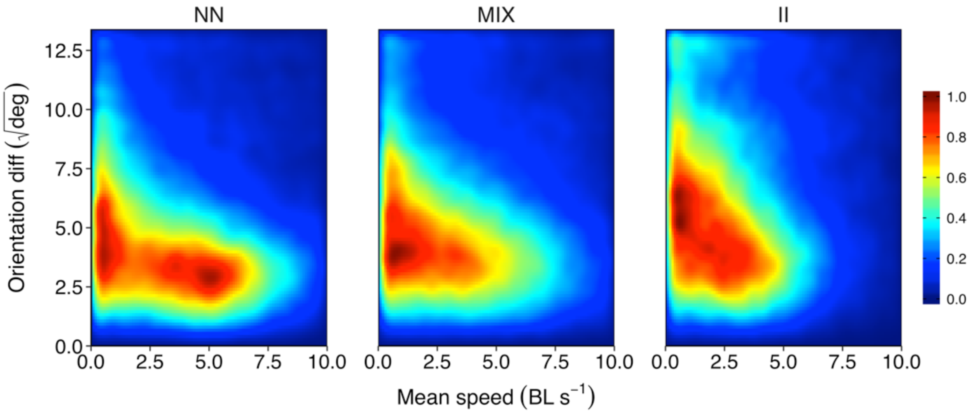


**Figure S10.** Heat maps showing the relationship between the mean speed of the fish in a pair and the difference in their orientation (square-root transformed) based on the full frame-by-frame dataset of the pair assay separately for pairs consisting of two non-infected fish (NN; $n$ = 17), of one infected and one non-infected fish (MIX; $n$ = 21), and two infected fish (II; $n$ = 21). Data is cropped to show the most relevant area only, respectively 95.2%, 95.1%, and 98.0% of the full parameter space. Colour scale is proportional to the densest bin of all plots.

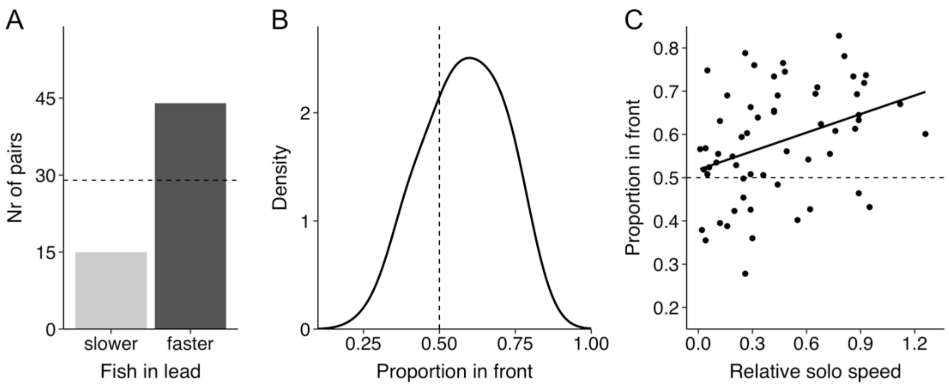


**Figure S11.** A) Number of pairs (*n* = 59) for which the fish in lead had the higher or lower solo speed ('faster' and 'slower' respectively). Dashed line indicates chance level. B) Density plot of the proportion of time the fish with the higher solo speed was the front-most fish during the trial. C) Scatterplot showing the relationship between fish' relative solo speed towards their partner and the proportion of time spent in front (*n* = 59 pairs).

### Appendix C: Agent-based modelling simulations

#### Individual-based modelling methods

*Overview*

We use a modified version of the zonal model [4,5], which uses persistent-turning-walker dynamics [6,7] to represent how individuals turn in response to social neighbours. The model represents each agent with an Ornstein-Uhlenbeck stochastic processes for changes in angular velocity $\omega$, and includes social interactions via a "social torque" based on the zonal model [3–5]. The speed ($s$) of each agent also is described by an Ornstein-Uhlenbeck stochastic process [7]. Motivated by the observed speed distributions of the non-infected and infected fish, we further extend this framework to incorporate stop-go motion, where individuals alternate between regular ("go") and low speed ("stop") states, with switches between states determined by both a preferred state dwelling time and social state synchronization coupling. The simulation model allows us to ask how coherence, alignment, and leadership are influenced by multiple different factors: 1) turning responsiveness, 2) turning noise, 3) movement speed, 4) preferred dwelling times in go/stop states.

*Turning dynamics*

Each individual in the simulation is characterized by the following time-dependent variables: position vector $\mathbf{r}_i = (x_i, y_i)$, unit velocity vector $\hat{\mathbf{v}}_i = (v_{i,x}, v_{i,y})$, speed $s_i$, and angular velocity $\omega_i$. The zonal model represents social interactions via three discrete 'zones' of repulsion, orientation, and attraction. We first give a simplified description of the zonal model specific to the pair simulations performed here. The three social zones of repulsion, orientation, and attraction are defined by the distance radii $r_R$, $r_O$, and $r_A$, respectively. For a focal individual $i$ and its neighbor $j$, let the distance between the two be $r_{ij} = |\mathbf{r}_j - \mathbf{r}_i|$. The preferred motion direction from the zonal model is determined as:

$$\hat{\mathbf{d}}(\mathbf{r}_i, \mathbf{r}_j, \hat{\mathbf{v}}_j) = \begin{cases} -\dfrac{\mathbf{r}_j - \mathbf{r}_i}{r_{ij}}, & 0 < r_{ij} \le r_R \\ \hat{\mathbf{v}}_j, & r_R < r_{ij} \le r_O \\ \dfrac{\mathbf{r}_j - \mathbf{r}_i}{r_{ij}}, & r_O < r_{ij} \le r_A \\ (0,0), & otherwise \end{cases}$$

This is used to calculated an effective 'social torque' according to

$$\Gamma_{\mathrm{ij}} = \left(\hat{\mathbf{v}}_i \times \hat{\mathbf{d}}_{ij}\right) \cdot \hat{z} = v_{i,x} d_{ij,y} - v_{i,y} d_{ij,x}$$

where $\hat{z}$ is a unit vector in the $z$-direction, and the notation $\hat{\mathbf{d}}_{ij} \equiv \hat{\mathbf{d}}(\mathbf{r}_i, \mathbf{r}_j, \hat{\mathbf{v}}_j)$ with components $(d_{ij,x}, d_{ij,y})$ is used. Individual $i$ is represented as a persistent turning walker via a stochastic differential equation for changes in angular velocity with time:

$$d\omega_i = \frac{1}{\tau}\left(\alpha_i\, \Gamma_{ij} - \omega_i\right)dt + \frac{\sigma}{\sqrt{\tau}} dW(t) \tag{1}$$

where $\tau$ is the angular velocity autocorrelation time, $\alpha_i$ is the social coupling strength (hereafter referred to as 'turning responsiveness'), $\sigma$ is the noise amplitude, and $W(t)$ is a Wiener noise process. We scale the amplitude of the noise process by the inverse square root of the autocorrelation time, so that in the absence of social interactions, the steady state variance of angular velocity is simply $\sigma^2/2$; with this formulation, the values of both $\alpha_i$ and $\sigma$ have units of radians/sec. Using Eq. 1, the agent's turning behaviour is determined by two factors: social interactions are set by the turning responsiveness parameter $\alpha_i$, and random turning is set by the noise parameter $\sigma$. In the simulations, we vary the turning responsiveness by either assigning the same value to each individual, i.e. by setting $\alpha = \alpha_i = \alpha_j$, or by assigning different values of $\alpha_i$and $\alpha_j$.

*Speed dynamics and stop-go model*

Similar to the angular velocity, the speed of each individual is also represented via an Ornstein-Uhlenbeck stochastic process:

$$ds_i = \frac{1}{\tau_s}(g_i\mu_i - s_i)dt + \frac{g_i\sigma_s}{\sqrt{\tau_s}} dW(t). \tag{2}$$

where $\tau_s$ is the speed autocorrelation time, $\mu_i$ is the average (or preferred) speed, $\sigma_s$ is the noise amplitude, $W(t)$ is a Wiener noise process, and $g_i = g_i(t)$ is the time-dependent stop-go speed multiplier. To simulate Eq. 2, we impose an additional rule to restrict the speed to non-negative values, i.e. at each time step we use the update rule $s_i = \max(s_i, 0)$. As in Eq. 1, we scale the amplitude of the noise process by the inverse square root of the autocorrelation time, so that when in the go-state ($g_i = 1$), the steady state variance of speed is simply $\sigma_s^2/2$.

The continuous motion model is represented by $g_i = 1$. With this, the agent's speed distribution is a truncated Gaussian distribution defined by mean and variance parameters $\mu_i$ and $\sigma_s^2/2$. The actual mean and variance will vary slightly from these values because of the imposed condition that speed is nonzero, but this difference is small for the parameter values used in the simulations here. For the continuous motion model, because the noise added to speed is uncorrelated between the agents, a small value of $\sigma_s$ does not affect the simulated trends with respect to turning behaviour. We therefore used $\sigma_s = 0.2$ for all simulations, both with the continuous motion and the stop-go model, because this value represents more realistic motion compared to using a strictly constant speed.

To represent stop-go motion, we define an additional rule to represent switching from a "go" state, where $g_i = 1$, to a "stop" state (or, more accurately, a "slow" state), where $g_i \ll 1$. To represent these changes, each agent has an accumulator variable $\eta_i$, and state switches occur when $\eta_i$ reaches a threshold of 1. The accumulator variable has dynamics described by

(3)

$$d\eta_i = \frac{1}{T(\text{state})}\left(1 - \alpha_g m(g_i, g_j)\right) dt + \sigma_g dW(t)$$

where $T(\text{state})$ is the switching time scale for the current state (i.e. either stop or go), $\alpha_g$ is a social weight for stop-go state coupling, $m(g_i, g_j)$ is the social coupling function, $W(t)$ is a Wiener noise process, and $\sigma_g$ is the noise amplitude. In addition to Eq. 3, we add an additional rule to prevent an agent from getting stuck in the stop or go states, by enforcing that $y_i$ must stay positive by applying the update rule $\eta_i = max(\eta_i, 0)$ at each time step. The social coupling function $m(g_i, g_j)$ takes a nonzero value if the agents are within the maximum social interaction radius, taking a value 1 or -1 depending on whether the agents are in the same or different states, i.e.

$$m(g_i, g_j) = \begin{cases} 1, & (r_{ij} \le r_A) \text{ and } (i \text{ and } j \text{ in same states}) \\ -1, & (r_{ij} \le r_A) \text{ and } (i \text{ and } j \text{ in different states}) \\ 0, & \text{otherwise} \end{cases}$$

The stop-go speed variable takes on value of

$$g_i = \begin{cases} 1, & \text{go state} \\ f_g/\mu_i, & \text{stop state} \end{cases}$$

where the scaling in the stop state is done so that if agents have different preferred speeds in the go-state, their effective preferred speed in the stop state is the same (i.e. the average speed in the go state for agent $i$ is $\mu_i$, while the average speed in the stop state is $f_g$ for both agents).

To understand how Eq. 3 leads to coupled state changes for a pair of individuals, first consider the case of zero noise and zero social coupling, i.e. $\sigma_g = \alpha_g = 0$. In this case, the agent switches its state every $T$ seconds. With noise ($\sigma_g \neq 0$), there will be a distribution of the times spent in each state, and for small values $\sigma_g$, the distribution will be approximately Gaussian with a mean of $T$ (for larger values the mean will be less than $T$ due to the statistics of the first-passage problem with the Ornstein-Uhlenbeck process, as well as due to additional rule we use that $\eta_i$ must stay positive). Now consider zero noise but a social coupling of $\alpha_g = 1$. With this value, if the agents are in the same state and within the attraction zone, then Eq. 3 yields $d\eta_i = 0$, i.e. $\eta_i$ will not change and the agents will always stay in their current state. However, if the agents are in different states and within the attraction zone, they switch states faster, in this case a switching time of $T/2$. Thus, for values of $0 < \alpha_g < 1$, agents will tend to synchronize states, because being in the same state as the other agent will increase the time spent in the current state, while being in the opposite state will decrease the time spent in the current state. See figure S13 for an example of the time trace of the accumulator variable $\eta_i(t)$ along with the speed state variable $g_i(t)$ for two coupled agents.

Because of the coupled state changes in the stop-go model, the median speed of each agent is an emergent outcome of the interactions during the simulations, and thus unlike the continuous motion model it is no longer equal to the value of $\mu_i$ that is set for each agent. With the parameter values used in the simulations (tables S1 and S2), the median speed of each agent depends on the values of $T_{go}$ for each agent, and also weakly on the values of the social responsiveness. figure S13 shows how simulated outcomes for median speed difference depends on the difference in speed parameters for the agents ($\Delta\mu$). Because of this difference, we use the following method to compare the stop-go simulation results with the continuous model for different values of the median speed (figure S14D). First, multiple simulations were performed with different speed parameters and turning responsiveness parameters, and the median speed difference and front-back difference were calculated. Because the changes in results were smooth with respect to parameter changes, we then used an interpolating function to determine the model result for the specified median speed differences shown in figure S14D. For figure S14C, we used an interpolating function to resample results with equal spacing in order to facilitate a comparison with the continuous motion model results shown in figure S14A.

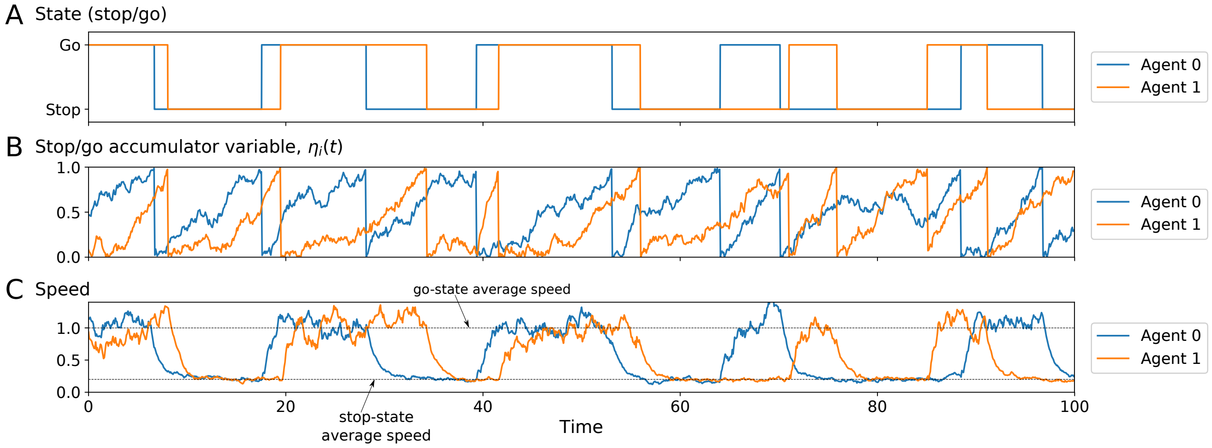


**Figure S12.** Example of the stop-go simulation model, showing synchronized changes in states. (A) Stop/go state for each agent. (B) Stop/go accumulator variable for each agent. The agent switches state when the accumulator variable reaches a threshold of 1; following this, $\eta_i$ resets to zero. (C) Speed for each agent during stop-go behavior. In this example, the average speed in the go-state is 1, and the average speed in the stop state is 0.2.

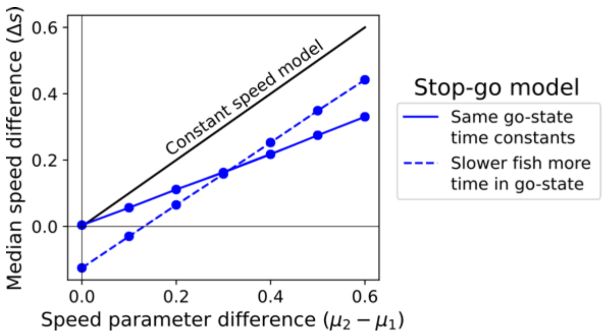


**Figure S13.** For the stop-go model, the simulation result for median speed difference (y-axis) is less than the speed parameter difference (x-axis), due to the coupling of the state changes. The median speed difference of the simulated agents also depends on the values of the go-state time constant. The solid blue line uses $T_{go} = 10$ for both agents, and the dashed blue line uses $T_{go} = 10$ for the slower agent and $T_{go} = 40$ for the slower agent. Both agents have social responsiveness values of $\alpha = 0.5$.

*Parameter values and simulation details*

The time and length scales in the simulation are set by the values of $\tau_s$ and $\mu\tau_s$, respectively, where $\mu = (\mu_1 + \mu_e)/2$ is the average speed parameter of the two agents. We choose $\tau_s = 1$, and $\mu = 1$. Thus, all time values are in units of $\tau_s$, and all distance values are in units of $\mu\tau_s$. Note that because the average pair speed defines the distance units, only the relative speed difference between agents, and not the absolute speed, is a meaningful quantity for comparison; we choose these definitions to simplify the number of parameters varied in the simulations.

The angular velocity autocorrelation time was chosen to be small compared to the speed autocorrelation time because this is what is seen in the data; we choose $\tau = 0.1\tau_s$. The values of $r_R$ and $r_O$ were chosen to correspond to the values used by [5]. However, there they had only a single social zone, which included both alignment and attraction. Here, we set the value of the alignment zone size to same as the social zone size from Ioannou et al. [5], and set a larger value for the attraction zone. Values of other parameters are listed in Table A1.

Simulations were performed by placing two individuals in a square arena of edge length 60, with periodic boundaries. In order to avoid any effects of initial conditions, we added an additional rule where all social interactions were set to zero for 4 $\tau_s$ after each time interval of 200 $\tau_s$. We ran simulations for a total time of $4 \times 10^5$, using a value of dt = 0.02. All simulations codes were written in Python 3.7.0 and are available on GitHub (https://git.io/Je6OT).

**Individual-based modelling results**

Group cohesion (lower IID) and alignment (lower absolute heading difference) were positively linked with a higher turning responsiveness for low to intermediate values of $\alpha$, and beyond these values there is little change (Figure S14A, C). In contrast, turning noise decreases group cohesion, but the effect depends on turning responsiveness; the spacing between agents with higher turning responsiveness ($\alpha = 0.5$) does not change much with higher noise values, but agents with lower turning responsiveness ($\alpha = 0.2$) have higher spacing when noise increases (Figure 14B). Turning noise also decreases group alignment, with a weaker effect for simulations with higher social responsiveness (Figure 14D). These results for group cohesion can be explained as follows: when turning responsiveness is sufficiently high compared to noise, the simulated agents move together and remain in each other's social interaction zones. When turning responsiveness is sufficiently low relative to noise, the agents sometimes move together and sometimes move apart. The use of discrete zones used in the model means that IID does not continue to decrease, but instead the median IID values remain just above the repulsion zone radius (see Table S1).

In terms of leadership, differences in speed strongly drive front/back positioning, with faster individuals being more in front (Figure S15). Also differences in social turning drive leadership, with the individual that has the lower turning responsiveness being more likely to lead. The stop-go model furthermore shows that differences in go-state dwelling time also have a small but consistent effect on leadership; when median speed and responsiveness are equal, the fish with high time spent in the go-state is more likely to lead (Figure S15C, D). Importantly, speed has a stronger effect on leadership than either turning responsiveness or go-state dwelling times and even in extreme cases, speed differences can overcome turning responsiveness differences such that the faster agent is more often in the front (Figure S15A, C; blues lines). With moderate speed differences, relatively large differences in turning responsiveness are needed to significantly affect leadership (Figure S15B, D). These results also show that using the more realistic stop-go model gives qualitatively similar results as the simpler continuous motion model.

Importantly, differences in turning responsiveness in our model could not only represent a difference in social tendency between two interacting individuals but also a difference in turning ability between two individuals with the same social tendency. Hence, the finding that pairs of agents with lower social responsiveness are less cohesive and aligned is in line with the finding that pairs of infected fish, which have lower turning rates, are less cohesive and aligned.

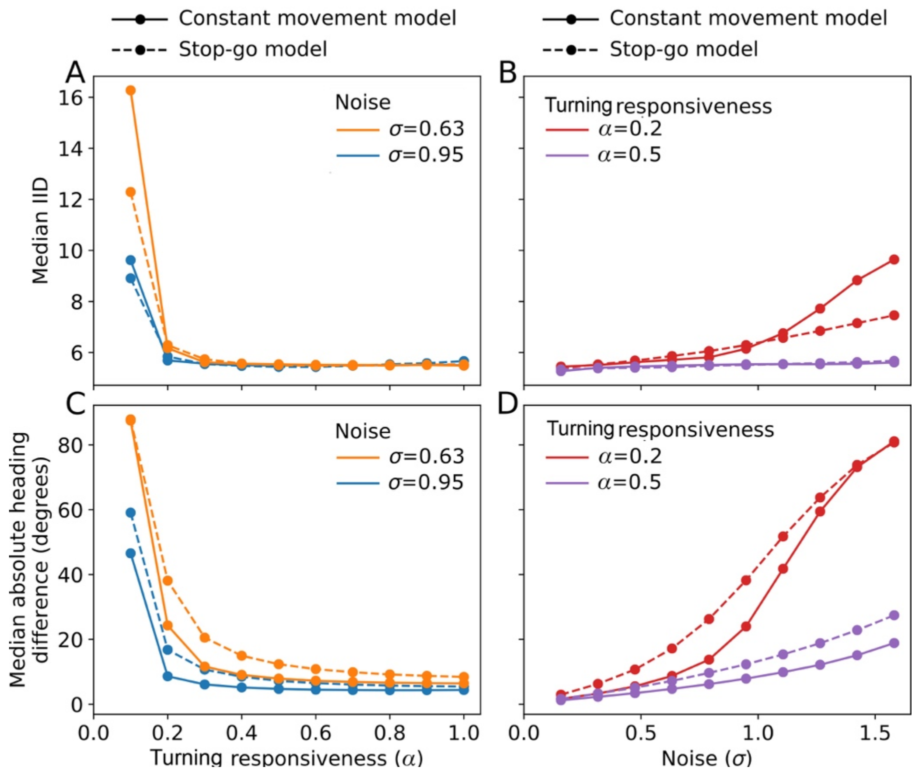


**Figure S14**. Plots of the agent-based model simulations of pairs of agents with the same median speeds but different values of turning responsiveness (A, C), and different values of turning noise (B, D), and their effect on group cohesion (A, B) and group alignment (C, D). Solid line indicates results of the continuous movement model, dashed line that of the stop-go model. The parameter values used for turning responsiveness and noise are specified above. Other parameter values are listed in Table S1. For the stop-go model, both agents had the same value of the go-state switching time constant: $T_{go} = 10$.

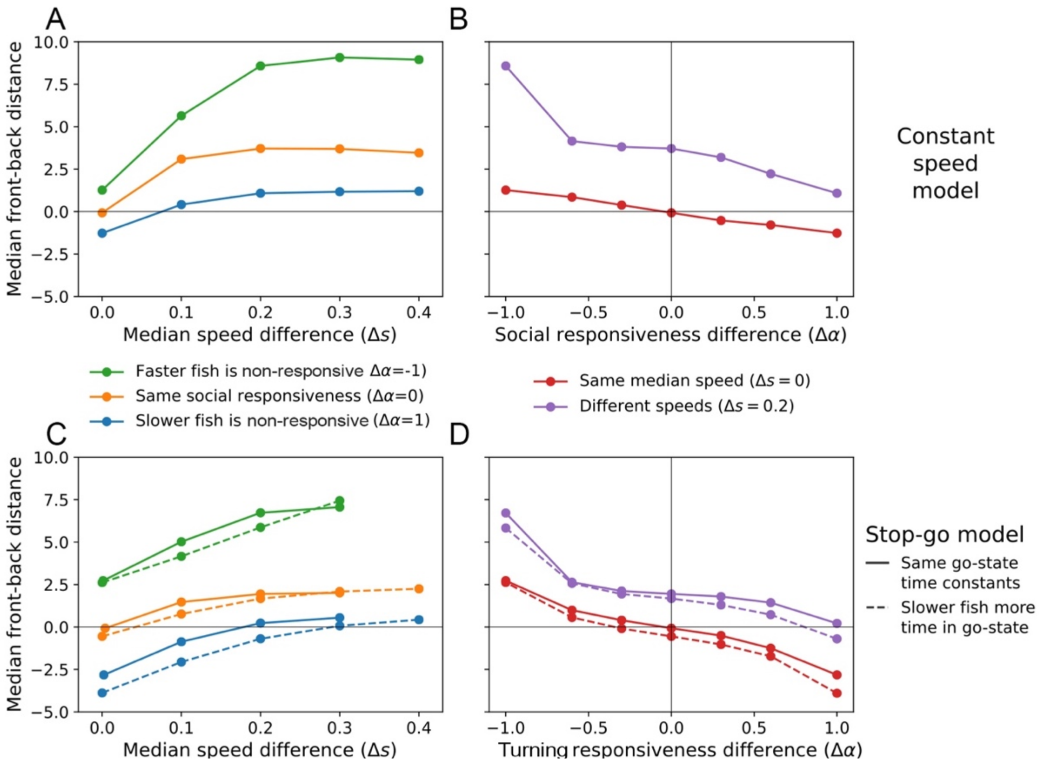


**Figure S15.** Agent-based modelling results for leadership in terms of front/back distance when individuals differ in speed and turning responsiveness, showing results for the continuous movement model (A, B) and for the stop-go model where agents have either the same ($T_{go} = 10$ for both; solid lines) or different values of the go state switching time constant ($T_{go} = 40$ for the slower fish, $T_{go} = 10$ for the faster fish; dashed lines) (C, D). Front/back distance is calculated with respect to the faster simulated fish; a positive value indicates the faster fish is the leader (i.e. spends more time in front). The turning responsiveness difference is also calculated with respect to the faster fish; thus $\Delta\alpha < 0$ means the faster fish is less responsive, and $\Delta\alpha > 0$ means the faster fish is more responsive. Panels (A, C) show leadership results as a function of median speed difference when agents have the same turning responsiveness of $\alpha = 0.5$ (orange lines), compared to the extreme cases where the faster fish is non-responsive ($\alpha = 0$) and the slower fish is responsive ($\alpha = 1$) (green lines), or where the slower fish is non-responsive and the faster fish is responsive (blue lines). Panels (B, D) show leadership results as a function of the turning responsiveness difference (mean social responsiveness of the pair is 0.5), when the agents have either the same (red) or different (purple) median speeds. See Tables S1 and S2 for other parameter values used.

**Table S1.** Fixed model parameter values used in the simulations.

| Parameter | Value | Description | Units |
|---|---|---|---|
| $\mu$ | 1 | Average speed parameter, $\mu = (\mu_1 + \mu_2)/2$ | Distance/time |
| $\tau_s$ | 1 | Speed autocorrelation time | Time unit |
| $\tau$ | 0.1 | Angular velocity autocorrelation time | $\tau_s$ |
| $\sigma_s$ | 0.2 | Speed noise amplitude | $\mu$ |
| $r_R$ | 3.657 | Repulsion zone | $\mu\tau_s$ (distance) |
| $r_O$ | 6.857 | Alignment/orientation zone | $\mu\tau_s$ (distance) |
| $r_A$ | 20 | Attraction zone | $\mu\tau_s$ (distance) |
| $\alpha_g$ | 0.8 | Social stop-go coupling | Non-dimensional |
| $f_g$ | 0.2 | Stop-state speed multiplier | Non-dimensional |
| $\sigma_g$ | 0.2 | Stop-go accumulator noise | $1/\sqrt{\tau_s}$ |
| $T_{stop}$ | 10 | Stop-state switching time | $\tau_s$ |

**Table S2.** Model parameters varied in the simulations. The description specifies which values are used in the two figures with the modelling results.

| Parameter | Values | Description | Units |
|---|---|---|---|
| $\sigma_i$ | 0.15-1.6,<br>0.95 | Angular velocity noise amplitude (Figure S13)<br>(Figure S14) | Rad/sec |
| $\Delta\mu = \mu_1 - \mu_2$ | 0-0.6 | Difference in preferred speed (Figure S14) | $\mu$ |
| $\alpha_i$ | 0-1 | Turning responsiveness | Rad/sec |
| $T_{go}$ | 10<br>10, 40 | Go-state switching time (Figure S13)<br>(Figure S14) | $\tau_s$ |
